# Iron Regulates CD4 T Cell Quiescence by Controlling TGF-β Production

**DOI:** 10.64898/2026.06.18.733147

**Authors:** Amber L. Siglin, Zelong Han, Afia Nkansah, WanJun Chen, Cheong-Hee Chang

## Abstract

Transforming growth factor-β (TGF-β) regulates CD4 T cell quiescence, activation, and regulatory T cell differentiation, but its role in T cell iron metabolism is poorly defined. Here, we investigated whether TGF-β regulates iron homeostasis and how iron overload alters TGF-β responsiveness. During T cell activation, TGF-β enhanced survival but markedly reduced proliferation. These effects were accompanied by decreased CD71 expression and cytosolic iron availability, as well as increased mitochondrial iron accumulation. Genetic deletion of TGFβR1 reversed these changes, demonstrating that TGF-β regulates CD4 T cell iron homeostasis through TGFβR1-dependent signaling.

Iron-overloaded CD4 T cells lacking the heme exporter FLVCR1 exhibit hypersensitivity to TGF-β, increased TGF-β secretion, and sustained TGFβR1 expression upon activation. Pharmacologic inhibition of TGFβR1restored proliferation, CD71 expression, and iron levels in FLVCR1-deficient cells. Although TGF-β selectively induced total and mitochondrial ROS levels in FLVCR1-deficient cells, antioxidant treatment or Nox2 inhibition did not rescue this phenotype, suggesting that ROS is associated with, but not sufficient to explain, TGF-β hypersensitivity. Acute FeSO_4_-induced iron overload partially recapitulated the phenotype of FLVCR1-deficient cells, although TGFβR1 expression and TGF-β production differed. Finally, regulatory T cells generated *in vitro* in the presence of TGF-β displayed reduced iron acquisition, and excess iron impaired FoxP3 induction. Together, this work identifies TGF-β as a context-dependent regulator of CD4 T cell iron homeostasis.

## INTRODUCTION

Iron is an essential micronutrient that supports immune cell survival, activation, proliferation, and effector function (1). Dietary iron is absorbed in the intestine, exported into the circulation, and transported in the bloodstream primarily by the glycoprotein transferrin (1). Cells acquire transferrin-bound iron through transferrin receptor 1, TFR1/CD71, which mediates endocytosis of iron-loaded transferrin (2). Once inside the cell, iron enters the labile iron pool, a metabolically accessible form of intracellular iron used for diverse cellular processes, including DNA synthesis, mitochondrial respiration, and enzymatic reactions (3). However, because of its redox activity, excess labile iron can promote oxidative stress and cellular damage. Therefore, intracellular iron levels must be tightly regulated. Excess iron can be stored in ferritin complexes or exported by ferroportin (4).

In addition to non-heme iron, heme metabolism provides additional mechanisms for maintaining iron balance. Heme represents a major intracellular iron-containing molecule and is required for essential metabolic processes, including mitochondrial oxidative phosphorylation. (5). Although *de novo* heme synthesis is a fundamental cellular process, heme can also be transported across membranes by specific transporters, including the heme importer FLVCR2 (6). Heme is degraded by heme oxygenase-1, HO-1, releasing free iron, biliverdin, and carbon monoxide (7). To prevent heme-associated toxicity, excess intracellular heme can be exported by the heme exporter FLVCR1 (8). Together, these pathways maintain iron and heme homeostasis and protect cells from iron-mediated oxidative injury.

Naïve CD4 T cells circulate through peripheral lymphoid tissues in a metabolically restrained state known as quiescence. This state is not passive; rather, it is actively maintained by extrinsic survival signals and low-level constitutive T cell receptor, TCR, engagement with self-peptide–MHC complexes, a process termed tonic signaling (9). In contrast, strong antigen-dependent TCR stimulation induces naïve CD4 T cells to exit quiescence and undergo activation, clonal expansion, and extensive metabolic reprogramming (10). Our group previously identified a physiological role for iron in maintaining naïve CD4 T cell quiescence using a mouse model with CD4 T cell-specific deficiency of FLVCR1 (11). Loss of FLVCR1 resulted in intracellular iron overload in naïve CD4 T cells. These iron-overloaded naïve CD4 T cells exhibited enhanced tonic signaling and spontaneous activation in vivo. Upon *in vitro* stimulation, FLVCR1-deficient CD4 T cells showed impaired activation responses, including reduced proliferation, decreased IL-2 production, and increased apoptosis and ferroptosis. In addition, stimulated iron-overloaded CD4 T cells displayed mitochondrial dysfunction and impaired metabolic fitness (12). These findings indicate that proper regulation of iron homeostasis is critical for maintaining the naïve CD4 T cell compartment and supporting productive T cell activation.

Transforming growth factor-β, TGF-β, is a pleiotropic cytokine that plays a central role in immune regulation (13). Among the TGF-β isoforms, TGF-β1 is the predominant isoform expressed in the immune system and regulates CD4 T cell survival, differentiation, tolerance, and effector function (13). TGF-β is secreted as a latent complex and requires activation, including release from latency-associated peptide before it can engage its receptors and initiate downstream signaling. Canonical TGF-β signaling begins when active TGF-β binds TGF-β receptor type II, TGFβR2, which recruits and phosphorylates TGF-β receptor type I, TGFβR1 (14). Activated TGFβR1 then phosphorylates receptor-associated SMAD proteins, primarily SMAD2 and SMAD3, which cooperate with SMAD4 to regulate transcriptional programs. In addition to canonical SMAD-dependent signaling, TGF-β receptors can also activate non-canonical pathways, including MAPK, ERK, and NF-κB-associated signaling cascades, thereby supporting context-dependent effects on cell survival, differentiation, and immune function (15).

Recent evidence suggests that TGFβR1 expression is an important regulator of naïve CD4 T cell quiescence (16). Following strong TCR stimulation, NF-κB activation suppresses TGFβR1 expression, limiting TGF-β responsiveness and allowing cells to exit quiescence. In contrast, low-level TCR stimulation, such as tonic signaling, is insufficient to downregulate TGFβR1, thereby permitting sustained TGF-β signaling and maintenance of the naïve quiescent state (16). These findings suggest that regulation of TGFβR1 expression may serve as a molecular checkpoint that distinguishes tonic signaling from productive antigen-driven activation.

TGF-β also plays a major role in CD4 T cell lineage differentiation, particularly in the induction of regulatory T cells, Treg (17). Treg are essential for maintaining immune tolerance, limiting immunopathology, and resolving immune responses after infection or inflammation (18). *In vitro*, TGF-β promotes the differentiation of induced Treg, iTreg, through induction of the lineage-defining transcription factor Foxp3 (17). Although TGF-β-dependent transcriptional and signaling pathways in Treg differentiation have been extensively studied, comparatively little is known about how iron homeostasis is regulated during iTreg differentiation or whether iron availability influences TGF-β-dependent Foxp3 induction. Understanding this relationship may provide insight into how metabolic and micronutrient pathways shape immune tolerance.

Together, prior work suggests that both iron homeostasis and TGF-β signaling are critical regulators of naïve CD4 T cell quiescence. However, whether these pathways are functionally connected remains unclear. In this study, we investigated the relationship between TGF-β signaling and iron metabolism in iron-overloaded naïve CD4 T cells upon stimulation, and iTreg-differentiated CD4 T cells. We show that TGF-β regulates CD4 T cell iron homeostasis through TGFβR1-dependent signaling, reducing CD71 expression and cytosolic iron availability while altering mitochondrial iron distribution. We further demonstrate that FLVCR1-deficient, iron-overloaded CD4 T cells exhibit enhanced TGF-β production, sustained TGFβR1 expression, and heightened sensitivity to TGF-β-mediated suppression of proliferation and iron acquisition.

Although reactive oxygen species (ROS) are increased in TGF-β-treated FLVCR1-deficient cells, antioxidant and Nox2-inhibition experiments suggest that ROS-associated pathways do not fully account for the TGF-β hypersensitive phenotype. Finally, we show that iTreg differentiation is associated with a distinct iron-homeostatic state and that excess iron impairs Foxp3 induction. Collectively, these findings identify TGF-β as a regulator of CD4 T cell iron homeostasis and suggest that iron availability shapes CD4 T cell responsiveness to TGF-β in a context-dependent manner.

## MATERIALS AND METHODS

### Mice

Male and female C57BL/6 mice ranging in age from 6-12 weeks were either bred in-house or purchased from Jackson Laboratories. FLVCR1-floxed mice were from Dr. Janis Abkowitz (U. of Washington). CD4^Cre^ mice were purchased from the Jackson Laboratories. FLVCR1 floxed mice were bred with CD4^Cre^ mice to produce FLVCR1^fl/fl^ (FLVCR1 WT) or FLVCR1^fl/fl^ CD4 ^Cre^ (FLVCR1 KO) mice. TGFβR1 ^fl/fl^ ER^Cre^ mice (16) were bred in-house. Mice were housed in specific pathogen free conditions. All animal experiments were performed in accordance with the Institutional Animal Care and Use Committee of the University of Michigan.

### Cell isolation, culture, and proliferation

Naïve CD4 T cells were enriched from C57BL/6 total splenocytes using enrichment kits according to the manufacturer’s instructions (Miltenyi Biotech and StemCell Tech). FLVCR1 WT and KO naïve CD4 T cells were sorted using ThermoFisher Bigfoot or Sony MA900. For cell proliferation measurements, naïve CD4 T cells were labeled with 2-5 uM CellTrace Violet (CTV, Invitrogen) in PBS with or without supplemented 1% BSA for 20 min at 37°C prior to stimulation. Naïve CD4 T cells were activated with plate-bound anti-CD3 (5 µg/ml) and soluble anti-CD28 (1 µg/mL) antibodies (eBioscience) in the presence or absence of recombinant TGF-β (Peprotech) for an indicated time in RPMI 1640 medium supplemented with 10% FBS and penicillin/streptomycin at 37°C.

TGFβR1 gene deletion was performed by culturing naïve cells from TGFβR1 ^fl/fl^ ER^Cre^ mice with 2 µM 4-hydroxytamoxifen

*In vitro* iron overload was induced in naïve CD4 T cells during activation by supplementing culture with 500 μM FeSO_4_ (Sigma Aldrich).

Induced regulatory T cells (iTreg) were differentiated from naïve CD4 T cells with a FoxP3 polarization cocktail containing 200 IU murine IL-2 (Peprotech), 2.5 ng/mL recombinant TGF-β (Peprotech), 1 µg/mL anti-IFNgamma, and 1 µg/mL anti-IL-4 (eBioscience) for 3 days. iTreg differentiation was confirmed by flow cytometry using the markers CD25 and FoxP3.

### Antioxidant Assays

Naïve CD4 T cells isolated from FLVCR1 WT and KO mice were treated with 5 mM N-acetyl cysteine (NAC, ThermoFisher) during 3 day stimulation. To inhibit Nox2 activity, cells were treated with 5 µM GSK2795039 (MedChemExpress LLC) during stimulation for 3 days.

### Flow cytometry assays

The fluorescently-conjugated antibodies used for surface and intracellular staining were: anti-mouse TCR-β (H57-597) PE-Cy7, anti-mouse CD4 (GK1.5) APC-Cy7, anti-mouse CD71 (R17217) FITC, anti-mouse CD25 (PC61.5) PE, anti-mouse CD44 (IM7) PerCP-Cy5.5, anti-mouse CD62L (MEL-14) BV605 (all from eBioscience). For Ferriportin staining, cells were fixed with 4% PFA in the dark at room temperature for 10 minutes. Fixed cells were incubated with a metal transporter protein antibody (rabbit anti-mouse MTP1/IREG1/Ferroprotein, Fpn) (Alpha Diagnostic) in flow cytometry buffer. Ferritin expression was measured by anti-mouse ferritin (EPR3004Y) (Abcam) staining in permeabilization buffer after fixation. Heme oxygenase-1 (HO-1) expression was measured by anti-mouse HO-1 (EP1391Y) (Abcam) in permealization buffer after fixation. Cells were labeled with Goat anti-rabbit IgG secondary antibody (A27034) (Invitrogen) before being subjected to flow cytometry analysis. iTreg were stained with anti-FoxP3 (FJK-16S)-APC using the FoxP3/transcription factor buffer kit per the manufacturer’s instructions (eBioscience).

Cells were acquired on a LSR Fortessa (BD Bioscience) and data was analyzed using FlowJo (TreeStar software ver. 10.5). Dead cells were excluded from the analysis based on Live/dead Fixable Aqua or Yellow (1 μg/mL) (Invitrogen) signal.

### Mitochondrial Function

Mitochondrial ROS (mitoROS) was measured using the dye MitoSOX (2.5 µM, Invitrogen). To measure mitochondrial iron, cells were labeled using MitoFerrogreen (5 µM, Dojindo). Labeling was done by incubating cells with the described dyes for 30 min at 37°C in RPMI 1640 media supplemented with 10% FBS and pen/strep. Cells were washed twice with flow cytometry buffer before being subjected to flow cytometry.

### RT-qPCR

Total RNA was isolated from both unstimulated and stimulated naïve CD4 T cells using the RNeasy Plus mini kit (Qiagen) according to the manufacturer’s instructions. cDNA was reverse transcribed, and RT-qPCR was performed using SYBR Green with Applied Biosystem’s 7500HT Sequence Detection System. Fold changes were calculated from Ct values using the ΔCt method. Expression of target genes was normalized to β-actin. Primers for β-actin (F: TTCGTTGCCGGTCCACA, R: ACCAGCGCAGCGATATCG), *Tgfb1* (F: TGATACGCCTGAGTGGCTGTCT, R: CACAAGAGCAGTGAGCGCTGAA), *Tgfbr1* (F: CAGAGGGCACCACCTTAAAA, R: AATGGTCCTGGCAATTGTTC), *Tfrc* (F: GAAGTCCAGTGTGGGAACAGGT, R: CAACCACTCAGTGGCACCAACA), *Nox2* (F: CCCAACTGGGATAACGAGTTCA, R: AGG GCC ACA CAG GAA AACG), and *il2* (F CCTGAGCAGGATGGAGAATTACA, R: TCCAGAACATGCCGCAGAG) were acquired from IDT.

### ELISA

Media supernatant was collected from stimulated naïve CD4 T cell cultures and centrifuged briefly to pellet cell debris. ELISA assays were completed in conjunction with the University of Michigan ELISA core. Secreted protein was calculated as ng/mL produced by 1,000,000 cells.

### Statistical analysis

All graphs were prepared using Prism software (Prism version 9; Graphpad Software, San Diego, CA). For comparison among multiple groups, the data was analyzed using a one-way ANOVA with a multi-comparison post-hoc test. For comparison between two groups, unpaired Student T-tests were used. P < 0.05 was considered statistically significant.

## RESULTS

### TGF-β Controls Iron Homeostasis in CD4 T Cells

We previously reported a role for TGF-β in maintaining naïve CD4 T cell quiescence (16). Additionally, we demonstrated that iron regulates naïve CD4 T cell survival by modulating tonic signaling (12). This association prompted us to investigate whether TGF-β maintains naïve CD4 T cell quiescence by regulating iron homeostasis. To address this question, we examined the response of naïve CD4 T cells to TGF-β during activation. Splenic naïve CD4 T cells were enriched from C57BL/6 mice, labeled with CellTrace Violet (CTV) and activated for 3 days in the presence or absence of recombinant mouse TGF-β. TGF-β-treated cells showed increased survival compared to controls, but their proliferative capacity was greatly reduced (Figure 1A).

**Figure 1.**
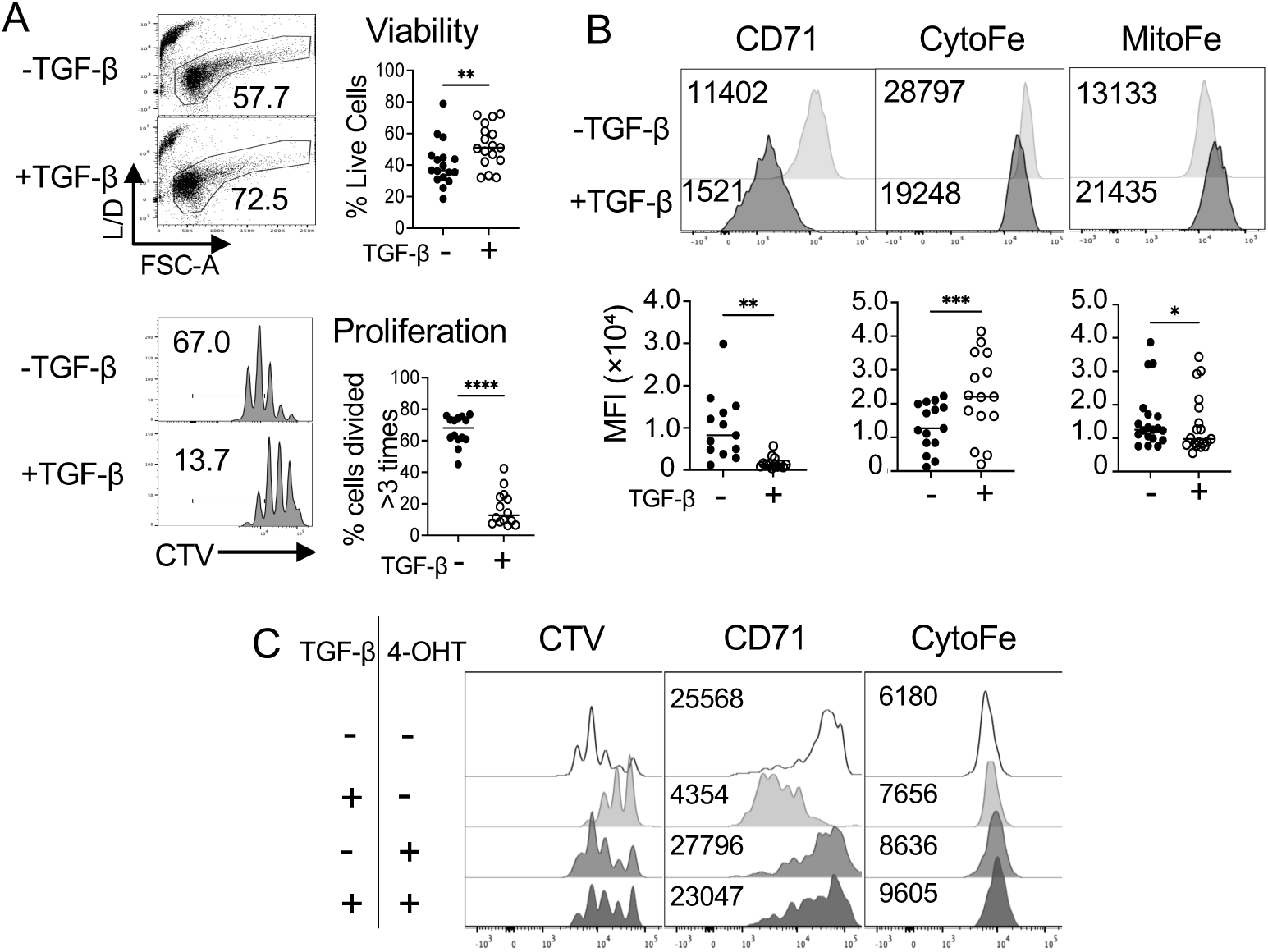
TGF-β Controls iron homeostasis in CD4 T cells. Naïve CD4 T cells were enriched from C57BL/6 splenocytes, labeled with 2-5 uM CellTrace Violet (CTV), and activated with 5 μg/mL plate-bound ⍺-CD3 and 1 μg/mL soluble ⍺-CD28 for 3 days in the presence or absence of 1 ng/mL TGF-β. **(A)** Cell viability assessed by live/dead staining, and proliferation by CTV dilution, are shown with summary graphs. n>14 **(B)** Representative histograms and summary graphs showing CD71 expression, iron levels in the cytosol (CytoFe) and the mitochondria (MitoFe). CytoFe and MitoFe levels were measured using FerroOrange and MitoFerroGreen, respectively, as described in Materials and Methods. n>13 **(C)** Naïve CD4 T cells were enriched from TGFβR1^fl/fl^ ER^Cre^ splenocytes. To induce deletion of *Tgfbr1*, 4-hydroxytamoxifen (4-OHT), (2 μM) was added during 3-day stimulation. Cells were cultured in the presence or absence of 1 ng/mL TGF-β. Proliferation, CD71 expression, and cytosolic iron levels were measured by flow cytometry after stimulation. Representative histograms are presented. n=2. Error bars represent the mean ± SEM. *P < 0.05, **P < 0.01, ***P < 0.001, ****P < 0.0001, ns: not significant.

To determine whether TGF-β regulates iron homeostasis, stimulated cells cultured with or without TGF-β were analyzed for CD71 expression, and iron levels in the cytosol (CytoFe) and the mitochondria (MitoFe). TGF-β treatment reduced CD71 expression, which was accompanied by decreased cytosolic iron levels and increased mitochondrial iron levels (Figure 1B). We next hypothesized that the effects of TGF-β on iron homeostasis are mediated through TGFβR1, a subunit of the TGF-β receptor complex (16). To test this, we used TGFβR1^fl/fl^ ER^Cre^ mice (16). Naïve CD4 T cells were stimulated with 4-hydroxytamoxifen (4-OHT) to induce TGFβR1 gene deletion. Deletion of TGFβR1 rescued TGF-β-mediated reduction in proliferation, CD71 expression, and cytosolic iron (Figure 1C). Together, these data support a role for TGFβR1 in the TGF-β-dependent regulation of iron homeostasis in CD4 T cells.

### Iron-Overloaded CD4 T Cells Exhibit Enhanced TGF-β Production and Sensitivity

We previously reported that CD4 T cells from mice lacking the heme exporter FLVCR1 exhibited elevated intracellular iron levels, resulting in impaired survival and proliferation, altered metabolism, and mitochondrial dysfunction (12). Moreover, these mice exhibit a markedly reduced naïve CD4 T cell compartment. Based on these observations, we investigated whether TGF-β contributes to the functional defects observed in iron-overloaded CD4 T cells. To test this, we used FLVCR1 wild type (FLVCR^fl/fl^) and FLVCR1 deficient mice (FLVCR1^fl/fl^-CD4^Cre^) as we have done previously (11,12). They are referred as WT and KO, respectively.

Naïve splenic CD4 T cells from WT and KO mice were sorted by flow cytometry, labeled with CellTrace Violet (CTV) and stimulated for 3 days in the presence or absence of TGF-β. Upon TGF-β treatment, KO cells exhibited a greater reduction in viability and proliferation than WT cells (Figure 2A), suggesting that iron-overloaded CD4 T cells are hypersensitive to TGF-β. Because 1 ng/mL TGF-β induced enhanced cell death in KO cells, we next performed a TGF-β titration. KO cells were sensitive to lower concentration of TGF-β compared to WT cells, further supporting heightened responsiveness to TGF-β in iron-overloaded cells (Figure 2B). We then examined parameters associated with iron homeostasis. TGF-β treatment reduced CD71 expression, which correlated with reduced cytosolic iron levels. In contrast to WT cells, KO cells did not exhibit enhanced mitochondrial iron accumulation upon TGF-β treatment (Figure 2C). These findings suggest that iron-overloaded CD4 T cells display altered iron redistribution in response to TGF-β.

**Figure 2.**
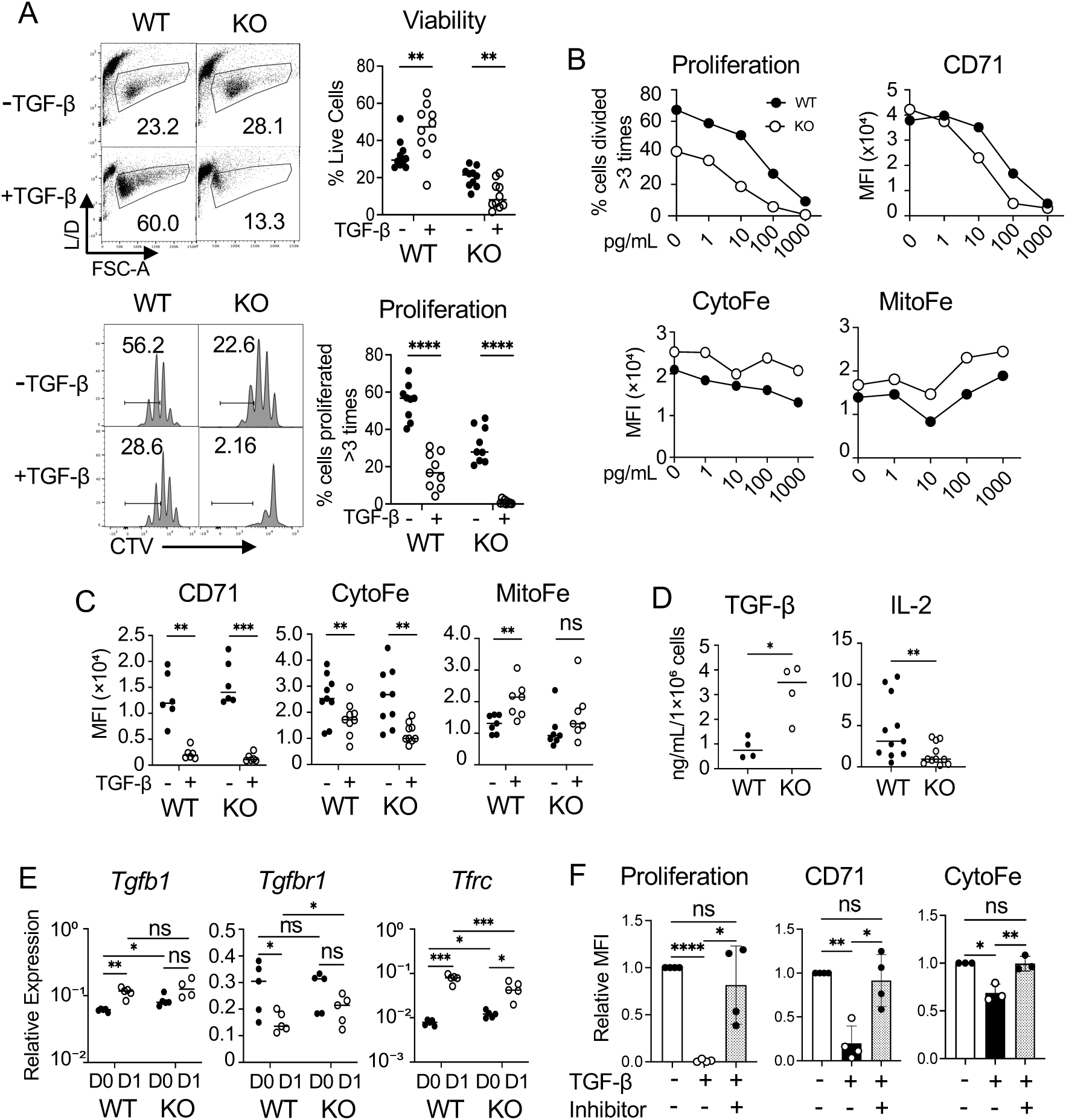
Iron-Overloaded CD4 T Cells Exhibit Enhanced TGF-β Production and Sensitivity. Naïve CD4 T cells from WT and KO mice were sorted by flow cytometry, labeled with CTV, and activated in the presence or absence of TGF-β. **(A)** Representative plots and histograms with summary graphs of viability and proliferation were shown. n>9 **(B)** WT and KO naïve CD4 T cells were stimulated for 3 days in the presence of increasing concentrations of TGF-β. Indicated parameters were measured by flow cytometry. **(C)** WT and KO naïve CD4 T cells were stimulated with or without TGF-β for 3 days, followed by measuring the levels of CD71 and iron. n>7 **(D)** WT and KO naïve CD4 T cells were stimulated for 3 days. Culture supernatants were collected and analyzed by ELISA. Total TGF-β was measured. n>4 **(E)** WT and KO naïve CD4 T cells were stimulated for 24 hours, and mRNA was prepared. RT-qPCR was performed to measure transcript levels of indicated genes. Relative expression was calculated using β-actin as the housekeeping gene. n=5 **(F)** Naïve KO CD4 T cells were treated with TGF-β in the presence of absence of TGFβR1 inhibitor SB-431542 (5 μM) throughout the 3 day stimulation. Proliferation, CD71 expression, and cytosolic and mitochondrial iron were measured. MFIs were normalized to the TGF-β-treated group without the inhibitor. n>4. Error bars represent the mean ± SEM. *P < 0.05, **P < 0.01, ***P < 0.001, ****P < 0.0001, ns: not significant.

Having observed hypersensitive responses of KO cells to TGF-β, we asked whether these cells produce higher levels of TGF-β than WT cells. We cultured WT and KO naïve CD4 T cells for 3 days and collected supernatants for ELISA. KO cells secreted higher levels of TGF-β compared to WT, whereas IL-2 levels were lower in KO than WT (figure 2D). Because KO cells secreted more TGF-β, we next examined whether TGF-β gene, *Tgfb1,* expression was also elevated. We compared mRNA levels between WT and KO cells before stimulation and after 24 hours of stimulation by RT-qPCR. Before stimulation, KO cells expressed higher levels of *Tgfb1* mRNA than WT. However, levels were comparable between WT and KO cells following 24 hours of stimulation. We also measured mRNA expression of *Tgfbr1*, which encodes TGFβR1. TGFβR1 expression has been reported to rapidly decrease upon stimulation, thereby limiting TGF-β signaling (16). Consistent with this, *Tgfbr1* expression was significantly reduced 24 hours after stimulation in WT cells. On the other hand, KO cells failed to downregulate *Tgfbr1* expression upon stimulation, suggesting that sustained TGFβR1 expression may permit persistent responsiveness to TGF-β. We also examined *Tfrc*, which encodes CD71. Before stimulation, *Tfrc* expression was higher in KO cells than WT cells. However, upon stimulation, KO cells failed to upregulate *Tfrc* to the same extent as WT cells (Figure 2E).

Our observation of sustained TGFβR1 expression in activated KO cells led us to hypothesize that inhibiting TGFβR1 signaling would restore proliferation and iron homeostasis in TGF-β-treated KO cells. To test this, we stimulated naïve KO CD4 T cells with or without SB-431542, a pharmacological inhibitor that targets the ATP binding domains of TGFβR1 and blocks downstream SMAD2/3 signaling (19). KO cells treated with the inhibitor exhibited improved proliferation, restored CD71 expression, and normalized intracellular iron levels toward control levels (Figure 2F). Together, this data suggests that heightened sensitivity to TGF-β in activated KO CD4 T cells is driven, at least in part, by sustained TGFβR1 expression.

### TGF-β Induces ROS Accumulation in Iron-Overloaded CD4 T Cells

Reactive oxygen species, ROS, can promote TGF-β production in various cell types, including T cells (20). Therefore, elevated TGF-β production in KO cells may be associated with increased ROS generation, potentially creating a detrimental negative feedback loop that impairs cell survival and proliferation. We first measured intracellular ROS and mitochondrial ROS in stimulated WT and KO cells cultured with or without TGF-β. In the absence of exogenous TGF-β, WT and KO cells generated comparable levels of total intracellular ROS and mitochondrial ROS following stimulation (figure 3A). However, when cells were treated with TGF-β, KO cells induced total and mitochondrial ROS, whereas WT cells did not (Figure 3A). These data suggest that iron-overloaded CD4 T cells are uniquely susceptible to TGF-β-induced oxidative stress.

**Figure 3.**
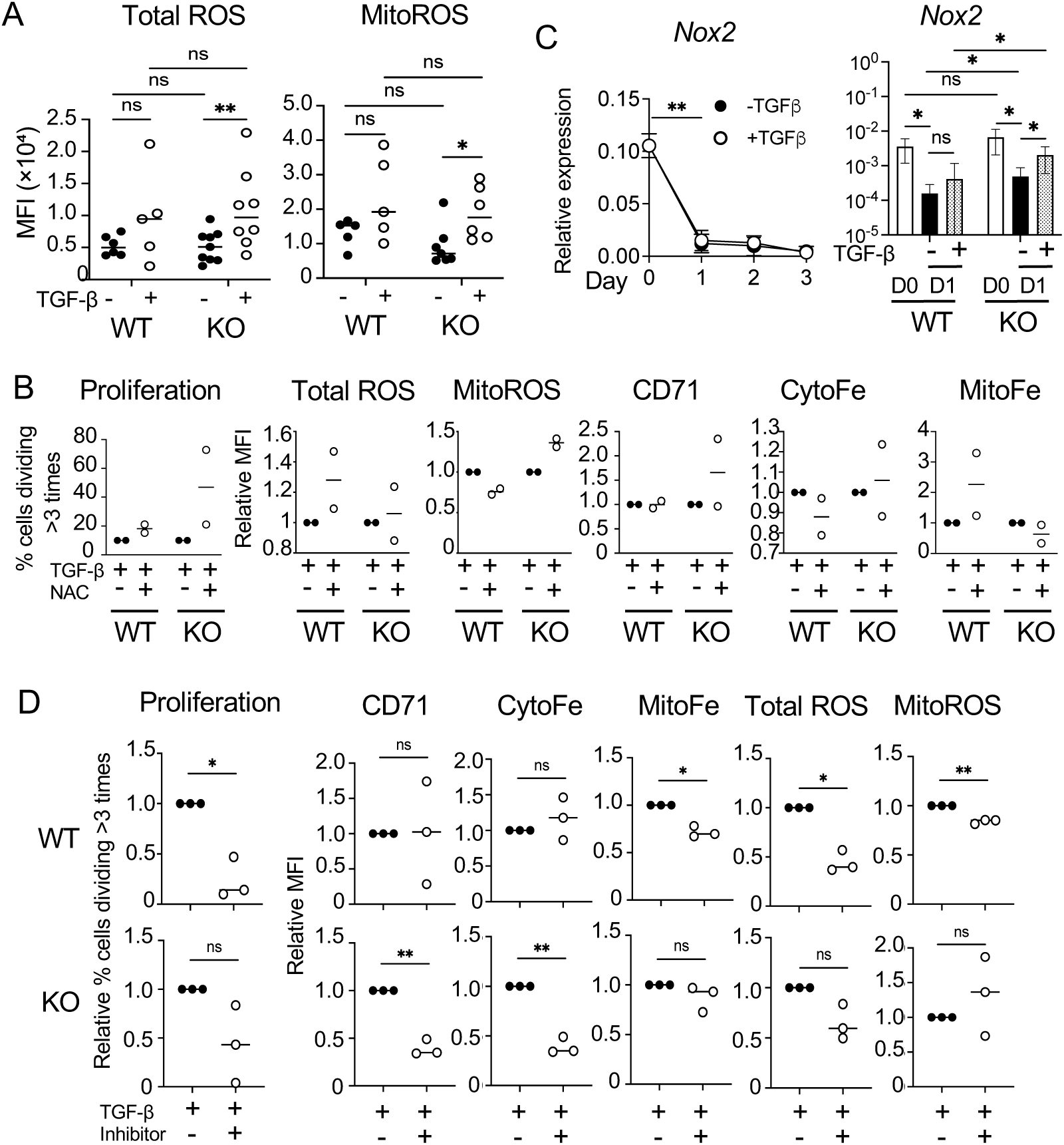
TGF-β Induces ROS Accumulation in Iron-Overloaded CD4 T Cells. Naïve CD4 T cells were sorted from WT and KO mice and stimulated in the presence or absence of TGFβ. **(A)** Summary graphs showing total intracellular ROS, Total ROS, and mitochondrial ROS, MitoROS, levels in activated WT and KO cells. Total ROS and MitoROS were measured using DCFDA and MitoSOX, respectively, as described in the Materials and Methods. n > 5. **(B)** WT and KO naïve CD4 T cells were stimulated with or without N-acetylcysteine (NAC, 5 mM) in the presence of TGF-β (1 ng/mL). Proliferation, total ROS, mitochondrial ROS, CD71, cytosolic iron, and mitochondrial iron were measured. MFIs were normalized to the TGF-ý-treated without the inhibitor. n=2 **(C)** Naïve CD4 T cells enriched from C57BL/6 spleens were stimulated in the presence or absence of 1 ng/mL TGF-β. *Nox2* mRNA levels were measured by RT-qPCR on days 1, 2, and 3 after stimulation in the presence or absence of TGF-β, left panel. n = 3. *Nox2* mRNA expression was also compared between WT and KO naïve CD4 T cells at day 0 and day 1 after stimulation with or without TGF-β, right panel. n > 4. **(D)** Naïve WT and KO cells were treated with 1 ng/mL or 30 pg/mL TGF-β, respectively, in the presence or absence of the Nox2 inhibitor GSK2795309 (5 μM) throughout 3 day stimulation. Proliferation, CD71, cytosolic and mitochondrial iron, and total and mitochondrial ROS levels were measured. MFIs were normalized to the TGF-β-treated group without the inhibitor. n=3. Error bars represent the mean ± SEM. *P < 0.05, **P < 0.01, ns: not significant.

Next, we asked whether elevated ROS contributes to the heightened TGF-β responseless in KO cells. To test this, cells were stimulated with or without the antioxidant N-Acetyl Cysteine (NAC) in the presence of TGF-β. NAC treatment in the presence of TGF-β failed to reduce ROS levels in WT or KO cells, and did not restore the other parameters tested, including proliferation and iron-associated phenotypes (Figure 3B). These results suggest that relevant ROS species or subcellular source may not be effectively targeted by NAC under these conditions.

Oxidative stress can activate transcriptional programs that regulates TGF-β gene expression (21). Because we observed elevated mitochondrial ROS in KO cells activated in the presence of TGF-β, we examined whether KO cells have altered expression of ROS-generating enzymes. We focused on the NADPH oxidase family, a major source of ROS in immune cells, including CD4 T cells (22). In particular, we examined Nox2, which is expressed in T cells and has been implicated in mitochondrial superoxide production through activation of reverse electron transfer (23). We found that over the course of 3 days of stimulation, *Nox2* transcript levels decreased within the first 24 hours and remained low throughout stimulation (Figure 3C, left panel). We then compared *Nox2* gene expression between WT and KO cells. *Nox2* mRNA expression was similar in unstimulated WT and KO cells (Figure 3C, right panel, compare D0). Upon stimulation, however, KO cells expressed higher levels of *Nox2* mRNA compared to WT cells, particularly following TGF-b treatment (Figure 3C, right panel). These findings suggest that ROS production in KO cells may be associated, at least in part, with increased *Nox2* expression.

We then hypothesized that Nox2-dependent ROS generation contributes to the elevated TGF-β production, reduced proliferation, and impaired iron homeostasis observed in KO cells. To test this, we stimulated cells with TGF-β in the presence or absence of GSK2795309, a pharmacologic inhibitor of Nox2 activity (24). For these experiments, KO cells were treated with 30 pg/mL of TGF-β to account for their increased TGF-β production and heightened TGF-β sensitivity. This concentration was chosen based on the titration data in Figure 2B, where 30 pg/mL TGF-β in KO cells showed effects comparable to those with 1 ng/mL TGF-β in WT cells. Unexpectedly, combined treatment with GSK2795309 and TGF-β further reduced proliferation in both WT and KO cells. However, WT and KO cells differed in their iron-associated responses. KO cells showed decreased CD71 expression and reduced cytosolic iron, whereas WT cells did not show measurable changes in these parameters. Mitochondrial iron and ROS levels were decreased in WT cells but not in KO cells (Figure 3D). These data suggest that Nox2 activity might be required to support optimal T cell activation and iron acquisition.

### *In Vitro* Iron Overload Partially Mimics TGF-β Hypersensitivity

We next asked whether the hypersensitivity of KO cells to TGF-β could be modeled using an *in vitro* iron-overloaded system. To induce iron overload, naïve CD4 T cells were activated in culture media supplemented with ferrous sulfate, FeSO_4_. Similar to KO cells, FeSO_4_-treated cells exhibited reduced viability and proliferation in response to TGF-β compared to untreated controls (figure 4A). These findings suggest that increased iron availability is sufficient to enhance TGF-β-associated defects in CD4 T cell survival and proliferation.

**Figure 4.**
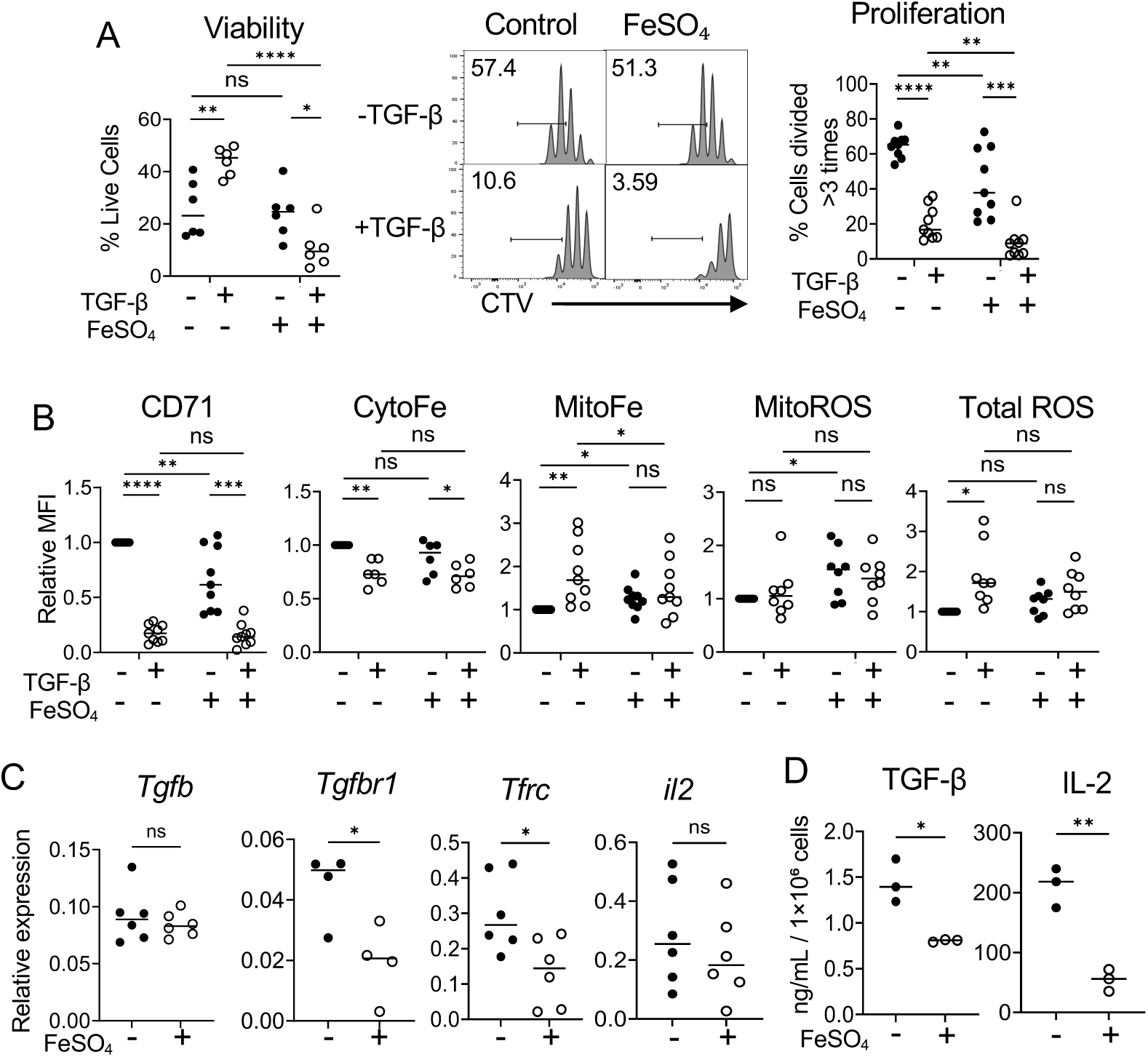
*In Vitro* Iron Overload Partially Mimics TGF-β Hypersensitivity. **(A,B)** Naïve CD4 T cells from C57BL/6 mice were stimulated for 3 days in the presence of TGF-β, 500 μM FeSO_4_, or TGF-β together with FeSO_4_. Representative histograms and summary graphs show viability and proliferation (**A**), and the levels of cytosolic iron, CD71, mitochondrial iron levels, and total and mitochondrial ROS level (**B**). n>6 **(C)** Naïve CD4 T cells were stimulated with or without FeSO_4_ for 24 hours. mRNA levels of indicated genes were measured by RT-qPCR. n>4 **(D)** Culture supernatants were collected after 3 days of stimulation and subjected to ELISA. n=3. Error bars represent the mean ± SEM. *P < 0.05, **P < 0.01, ***P < 0.001, ****P < 0.0001, ns: not significant.

Next, we studied the effect of FeSO_4_ on iron homeostasis. In the absence of TGF-β, cells activated in the presence of FeSO_4_ showed significantly reduced CD71 expression compared with untreated controls (Figure 4B). However, cytosolic iron levels remained comparable, whereas mitochondrial iron levels increased (Figure 4B). Similarly, mitochondrial ROS was induced while maintaining total ROS. TGF-β treatment of FeSO_4_ -induced iron-overloaded cells further reduced CD71 expression and cytosolic iron levels. However, TGF-β had little effect on mitochondrial iron and ROS levels (figure 4B). We then assessed mRNA expression of genes involved in iron homeostasis and TGF-β responsiveness. Consistent with the phenotype of KO cells, FeSO_4_ -induced iron overload reduces *Tfrc* expression. However, unlike KO cells, which maintained elevated *Tgfbr1* expression following activation, FeSO_4_-treated cells expressed lower levels of *Tgfbr1* mRNA compared with untreated controls. *Tgfb1* expression was comparable between groups (Figure 4C). The reduction in *Tgfbr1* expression was accompanied by reduced secretion of both TGF-β and IL-2 (Figure 4D).

Together, these findings indicate that *in vitro* iron overload partially recapitulates the TGF-β hypersensitivity observed in KO cells, particularly with respect to reduced viability, impaired proliferation, diminished CD71 expression, and altered intracellular iron distribution. However, the differences in *Tgfbr1* expression and TGF-β production suggest that the relationship between iron homeostasis and TGF-β responsiveness is context-dependent and may vary according to the mechanism by which iron overload is induced.

### Regulatory T Cells Differentiated with TGF-β *In Vitro* Show Distinct Iron Homeostasis

TGF-β plays a major role in immune tolerance and is a key factor required for the differentiation of induced regulatory T cells, iTreg. Given our finding that TGF-β regulates iron homeostasis in activated CD4 T cells, we next investigated iron metabolism during iTreg differentiation. Naïve CD4 T cells from C57BL/6 mice were stimulated under conventional T cell, Tconv, or iTreg differentiation conditions for 3 days. As expected, iTreg expressed high levels of FoxP3 and had reduced proliferation compared to Tconv cells (Figure 5A). Analysis of iron associated parameters revealed that iTreg displayed a distinct iron profile characterized by reduced iron availability compared to Tconv cells. Specifically, iTreg exhibited lower total iron levels, reduced CD71 expression, and decreased expression of ferritin, heme oxygenase-1, HO-1, and ferroportin compared to Tconv cells. Cytosolic iron levels were also lower in iTreg than in Tconv cells. However, mitochondrial iron and mitochondrial ROS levels were largely comparable between iTreg and Tconv cells (Figure 5B). These data suggest that iTreg differentiation is associated with reduced cytosolic iron availability and decreased expression of multiple iron-regulatory proteins.

**Figure 5.**
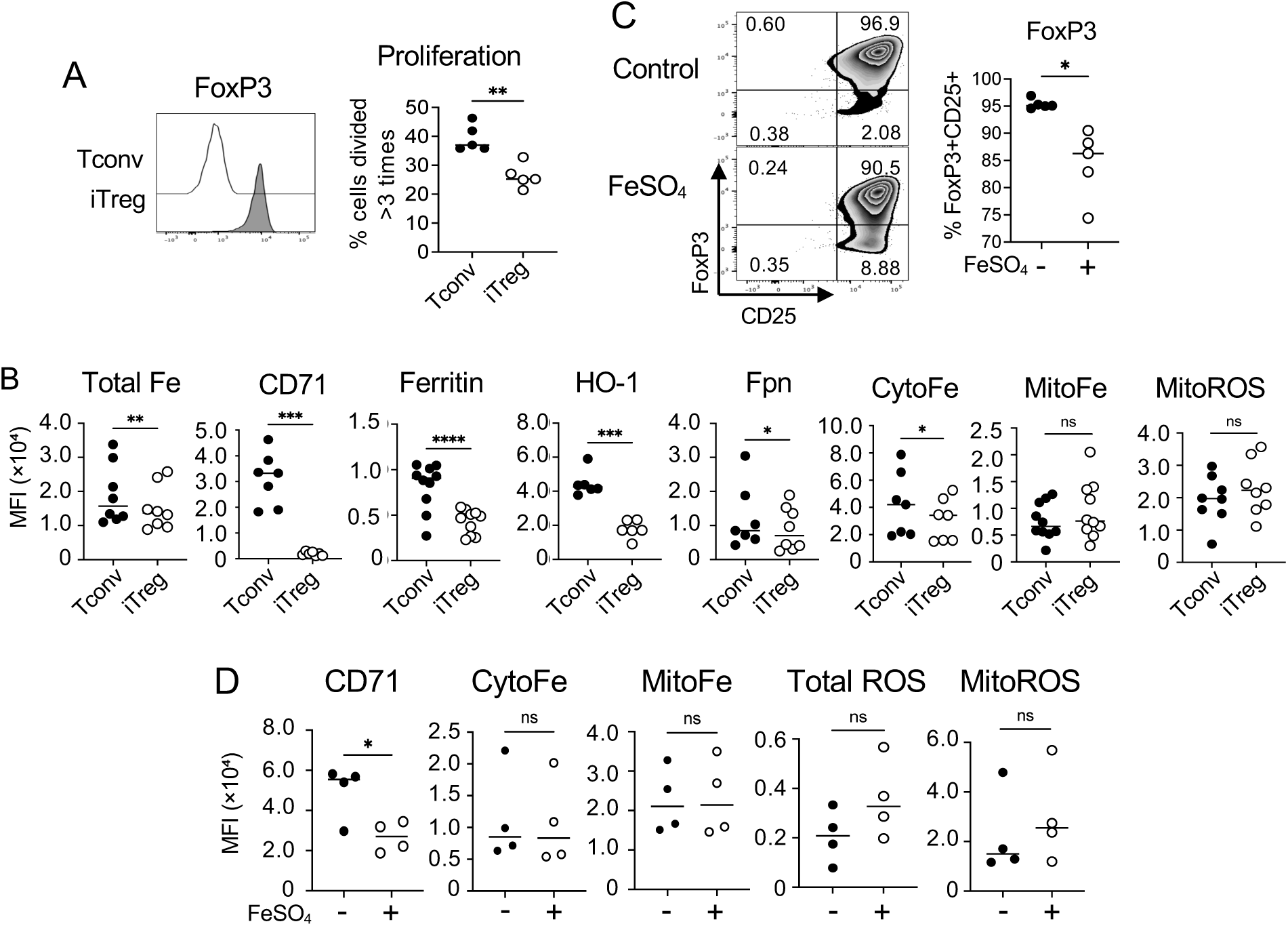
Regulatory T Cells Differentiated with TGF-β *In Vitro* Show Distinct Iron Homeostasis. Naïve CD4 T cells were enriched from C57BL/6 spleens and were differentiated into conventional T cells (Tconv) or induced regulatory T cells (iTreg) for 3 days. **(A)** Representative histogram of FoxP3 expression in Tconv and iTregs (left panel) and a summary graph of proliferation (right panel). n=5. **(B)** Tconv and iTreg were prepared as in (A), and levels of indicated parameters were compared. Total iron (total Fe) was measured by Calcein-AM as described in the Materials and Methods. n>6 **(C,D)** Naïve CD4 T cells were differneitated to iTreg with or without 500 μM FeSO_4_. Expression of FoxP3 with CD25 (**C**) and the levels of CD71, cytosolic and mitochondrial iron, total ROS, and mitochondrial ROS (**D**) are presented. n=4. Error bars represent the mean ± SEM. *P < 0.05, **P < 0.01, ***P < 0.001, ****P < 0.0001, ns: not significant.

To further investigate the effect of iron overload on TGF-β-dependent iTreg differentiation, we generated iTreg in the presence of FeSO_4_. FeSO_4_ treatment reduced FoxP3 expression, suggesting that excess iron impairs iTreg differentiation (Figure 5C). Additionally, FeSO_4_-treated iTreg showed reduced CD71 expression. However, intracellular iron and ROS levels were largely unchanged, suggesting that iTreg may use distinct regulatory mechanisms to maintain iron balance under conditions of increased extracellular iron availability (Figure 5D). Together, these findings indicate that TGF-β-dependent iTreg differentiation is associated with a unique iron-homeostatic state characterized by reduced CD71 expression, decreased cytosolic iron availability, and limited changes in mitochondrial iron. Moreover, excess iron disrupts FoxP3 induction, suggesting that proper regulation of iron homeostasis is required for efficient iTreg differentiation.

## DISCUSSION

Our current study identifies TGF-β as a regulator of iron homeostasis in CD4 T cells and suggests that iron availability shapes TGF-β responsiveness. TGF-β is known to inhibit T cell proliferation, promote iTreg differentiation, and support naïve T cell quiescence through regulated TGFβR1 expression. In naïve CD4 T cells, low-level tonic TCR signaling promotes peripheral survival but is insufficient to downregulate TGFβR1, thereby sustaining TGF-β signaling and quiescence (16,25,26). Previous work from our laboratory showed that FLVCR1-deficient CD4 T cells exhibit enhanced tonic TCR signaling. Here, we extend these findings by showing that FLVCR1-deficient, iron-overloaded CD4 T cells also display heightened TGF-β responsiveness.

In this study, we used two approaches to model iron overload in CD4 T cells: genetic disruption of heme export through FLVCR1 deletion and acute extracellular iron supplementation with FeSO_4_. Both models showed increased sensitivity to TGF-β-associated suppression of CD4 T cell proliferation and altered iron homeostasis. However, these models differed in their effects on TGF-β production and TGFβR1 expression. FLVCR1-deficient cells exhibited increased TGF-β secretion and sustained TGFβR1 expression following activation. In contrast, FeSO_4_-treated cells showed reduced TGFβR1 expression and decreased TGF-β production. These differences suggest that chronic intracellular iron or heme dysregulation caused by FLVCR1 deficiency is not equivalent to acute extracellular iron loading.

FLVCR1-deficient cells may experience sustained iron and heme stress beginning during development or naïve T cell maintenance, leading to persistent changes in signaling, metabolism, and TGF-β responsiveness. By contrast, FeSO_4_-treated cells may retain compensatory mechanisms that limit intracellular iron accumulation, such as reducing CD71 expression, decreasing iron import, limiting TGF-β receptor expression, or altering cytokine production. In addition, FeSO_4_ supplementation represents transient iron exposure, whereas FLVCR1 deficiency results in persistent disruption of iron homeostasis. Thus, iron overload is not a uniform cellular state. The biological consequences of excess iron likely depend on the route of iron accumulation, duration of exposure, subcellular iron localization, and the cell’s capacity to regulate iron import, storage, utilization, and export.

ROS are known to modulate TGF-β production and signaling in several cell types, including T cells. In our study, TGF-β treatment selectively increased total and mitochondrial ROS in FLVCR1-deficient cells. In addition, FLVCR1-deficient cells expressed higher levels of Nox2, suggesting that Nox2 may contribute to elevated ROS production. However, pharmacologic inhibition of Nox2 did not rescue the TGF-β-induced defects observed in FLVCR1-deficient cells. Instead, Nox2 inhibition further impaired proliferation and reduced iron-associated parameters, including CD71 expression and intracellular iron levels. Similarly, antioxidant treatment with NAC did not restore proliferation or iron homeostasis in TGF-β-treated FLVCR1-deficient cells. These findings suggest that although ROS accumulation is associated with TGF-β hypersensitivity in FLVCR1-deficient cells, ROS alone is unlikely to drive the phenotype. Rather, ROS may represent one component of a broader metabolic and signaling disruption caused by altered iron and heme homeostasis. Moreover, Nox2-dependent ROS may be required for optimal T cell activation rather than functioning solely as a pathogenic mediator in iron-overloaded CD4 T cells.

TGF-β is also essential for iTreg differentiation. Our data show that iTreg exhibit a distinct iron-homeostatic profile compared with activated CD4 Tconv cells. iTreg displayed reduced CD71 expression, lower cytosolic iron availability, and decreased expression of several iron-regulatory proteins, including ferritin, HO-1, and ferroportin. In contrast, mitochondrial iron and mitochondrial ROS levels were relatively preserved. These findings suggest that iTreg differentiation is associated with reduced iron acquisition and altered iron utilization compared with Tconv cell activation. When iTreg were differentiated in the presence of FeSO_4_, Foxp3 expression was reduced, indicating that excess extracellular iron can impair TGF-β-dependent iTreg differentiation. However, FeSO_4_ treatment had limited effects on measured intracellular iron parameters in iTreg. This suggests that iTreg may possess mechanisms that restrict iron accumulation or maintain iron balance during differentiation. Collectively, these data indicate that proper regulation of iron homeostasis is important for efficient Foxp3 induction and that excessive iron availability may disrupt iTreg differentiation.

Altered iron metabolism has been implicated in autoimmune diseases such as systemic lupus erythematosus, SLE, where increased CD71 expression and disrupted endosomal recycling promote iron uptake, mitochondrial dysfunction, enhanced Th17 differentiation, and impaired Treg development (27). Defective immunosuppressive TGF-β production has also been reported in SLE (28). In our study, acute FeSO_4_-induced iron overload reduced TGFβR1 expression and TGF-β production, whereas FLVCR1 deficiency sustained TGFβR1 expression and increased TGF-β production. These findings suggest that distinct forms of iron dysregulation differentially affect TGF-β signaling and CD4 T cell fate, with potential relevance to immune dysfunction in disease.

This study has limitations. First, because iron homeostasis is closely linked to cellular metabolism, future studies are warranted to investigte how TGF-β alters metabolic programming in iron-overloaded CD4 T cells. In particular, mitochondrial respiration, glycolysis, metabolite availability, and redox balance should be examined in detail during activation in the presence of TGF-β. Second, although our data show altered TGFβR1 expression and TGF-β responsiveness, we were limited in our ability to directly assess downstream TGF-β signaling in FLVCR1-deficient cells. The restricted naïve CD4 T cell compartment in FLVCR1 KO mice resulted in low naïve cell yield, limiting analysis of signaling dynamics. Future studies using inducible FLVCR1 deletion may help overcome this limitation and allow more direct analysis of canonical and non-canonical TGF-β signaling pathways in iron over-loaded CD4 T cells.

In conclusion, our study identifies TGF-β as a regulator of iron homeostasis during CD4 T cell activation. Iron-overloaded CD4 T cells showed increased TGF-β production, sustained TGFβR1 expression, and heightened TGF-β sensitivity. Acute FeSO_4_-induced iron overload only partially reproduced this phenotype, indicating that the effects of iron dysregulation depend on context and mechanism. We also show that iTreg differentiation is associated with reduced iron acquisition and that excess iron impairs Foxp3 induction. Together, these findings reveal a link between iron metabolism and TGF-β signaling that may shape CD4 T cell quiescence, activation, and regulatory T cell differentiation.

## ACKNOWLEDGEMENTS

We thank Dr. Bethany Moore (University of Michigan) for her support and helpful discussion, Isabelle Rolina and Adrianne Kurutz for mouse work, and the University of Michigan’s Flow Cytometry Core and Immune Monitoring Shared Resource for sample analysis.

## DISCLOSURE

The authors declare no conflict of interest.

## AUTHOR CONTRIBUTIONS

CHC conceived and designed the project. Experiments were performed by ALS, ZH, and AN. The data were analyzed and interpreted by CHC, ALS, ZH, and AN. The manuscript was prepared by CHC, ALS, and ZH. WC provided indispensable intellectual input. All authors reviewed and edited the manuscript

## Notes

### Competing Interest Statement

The authors have declared no competing interest.

